# HASE: Framework for efficient high-dimensional association analyses

**DOI:** 10.1101/037382

**Authors:** G.V. Roshchupkin, H.H.H. Adams, M.W. Vernooij, A. Hofman, C.M. Van Duijn, M.A. Ikram, W.J. Niessen

## Abstract

Large-scale data collection and processing have facilitated scientific discoveries in fields such as genomics and imaging, but cross-investigations between multiple big datasets remain impractical. Computational requirements of high-dimensional association studies are often too demanding for individual sites. Additionally, the sheer size of intermediate results is unfit for collaborative settings where summary statistics are exchanged for meta-analyses. Here we introduce the HASE framework to perform high-dimensional association studies with dramatic reduction in both computational burden and storage requirements of intermediate results. We implemented a novel meta-analytical method that yields identical power as pooled analyses without the need of sharing individual participant data. The efficiency of the framework is illustrated by associating 9 million genetic variants with 1.5 million brain imaging voxels in three cohorts (total N=4,034) followed by meta-analysis, on a standard computational infrastructure. These experiments indicate that HASE facilitates high-dimensional association studies enabling large multicenter association studies for future discoveries.

## INTRODUCTION

Technological innovations have enabled the large-scale acquisition of biological information from human subjects. The emergence of these big datasets has resulted in various ’omics’ fields. Systematic and large-scale investigations of DNA sequence variations (genomics)^1^, gene expression (transcriptomics)^2^, proteins (proteomics)^3^, small molecule metabolites (metabolomics)^4^, and medical images (radiomics)^5^, among other data, lie at the basis of many recent biological insights. These analyses are typically unidimensional, i.e. studying only a single disease or trait of interest.

Although this approach has proven its scientific merit through many discoveries, jointly investigating multiple big datasets would allow for their full exploitation, as is increasingly recognized throughout the ’omics’ world. However, the high-dimensional nature of these analyses makes them challenging and often unfeasible in current research settings. Specifically, the computational requirements for analyzing high-dimensional data are far beyond the infrastructural capabilities for single sites. Furthermore, it is incompatible with the typical collaborative approach of distributed multi-site analyses followed by meta-analysis, since the amount of generated data at every site is too large to transfer.

Some studies have attempted to combine multiple big datasets^5,8–10^, but these methods generally rely on reducing the dimensionality or making assumptions to approximate the results, which leads to a loss of information.

Here we present the High-dimensional Association Studies with Efficiency (HASE) framework, which is capable of analyzing high-dimensional data at full resolution, yielding exact association statistics (i.e. no approximations), and requiring only standard computational facilities. Additionally, the major computational burden in collaborative efforts is shifted from the individual sites to the meta-analytical level while at the same time reducing the amount of data needed to be exchanged and preserving participant privacy. HASE thus removes the current computational and logistic barriers for single-and multi-center analyses of big data.

## RESULTS

### Overview of the methods

The methods are described in detail in the Online Methods. Essentially, HASE implements a high-throughput multiple linear regression algorithm that is computationally efficient when analyzing high-dimensional data of any quantitative trait. Prior to analysis, data are converted to an optimized storage format to reduce reading and writing time. Redundant calculations are removed and the high-dimensional operations are simplified into a set of matrix operations that are computationally inexpensive, thereby reducing overall computational overhead. While deriving summary statistics (e.g., beta coefficients, p-values) for every combination in the high-dimensional analysis would be computationally feasible at individual sites with our approach, it would be too large to share the intermediate results (>200GB per thousand phenotypes) in a multi-center setting. Therefore, extending from a recently proposed method, partial derivatives^17^ meta-analysis, we additionally developed a method that generates two relatively small datasets (e.g. 5GB for genetics data of 9 million variants and 20MB of thousand phenotypes for 4000 individuals) that are easily transferred and can subsequently be combined to calculate the full set of summary statistics, without making any approximation. This meta-analysis method additionally reduces computational overhead at individual sites by shifting the most expensive calculation to the central site. The total computational burden thus becomes even more efficient relative to conventional methods with additional sites. The HASE software is freely available from our website www.imagene.nl/HASE/.

### Comparison of complexity and speed

We compared the complexity and speed of HASE with a classical workflow, based on linear regression analyses with PLINK (version 1.9)**^11^ followed by meta-analysis with METAL^12^**; two of the most popular software packages for these tasks.

**Table 1** shows that HASE dramatically reduces the complexity for the single site analysis and data transfer stages. For conventional methods, the single site analysis and data transfer have a multiplicative complexity (dependent on the number of phenotypes and determinants), whereas this is only additive for HASE. Our approach requires 2×10^6^-fold less time on the single site stage and 3.500-fold less data to transfer for a high-dimensional association study. Additionally, the time for single site analysis does not increase significantly from analyzing a single phenotype to a million phenotypes **(Table 1).** This is due to the fact that speed is determined by the highest number of either the determinants or phenotypes. Therefore, in this case with nine million genetic variants, the complexity of ***O*(*n_i_n_p_*)** is the primary factor influencing the speed, whereas ***O* (*n_i_n_t_***) plays a secondary role.

**Table 1.** Comparison of complexity and speed between the HASE framework and a classical workflow.

| Stage | Complexity<sup>c</sup> |  | Time<sup>a,b</sup><br>(hours) |  |  |  |
| --- | --- | --- | --- | --- | --- | --- |
| | Classical workflow | HASE | $n_p=1$ | | $n_p=10^6$ | |
|  |  |  | Classical workflow | HASE | Classical workflow | HASE |
| Single site analysis | $O(n_i n_p n_t)$ | $\max( O(n_i n_p), O(n_i n_t) )$ | 2.46 | 0.63 | $2.46 \times 10^6$ | 0.70 |
| Data transfer | $O(n_p n_t)$ | $O(n_i n_p + n_i n_t)$ | 0.04 | 0.07<sup>*</sup> | $4 \times 10^4$ | 11.6 |
| Meta-Analysis | $O(n_p n_t)$ | $O(n_i n_p n_t)$ | 0.06 | 0.03 | $6 \times 10^4$ | $1.7 \times 10^3$ |

<sup>a</sup> Based on a model with three covariates and 9 million genetic variants, for a total of 4034 participants from three sites. For the classical workflow we used the PLINK software for single site analysis and METAL for the meta-analysis.
<sup>b</sup> For single site analysis and meta-analysis the time is given in CPU hours; for the data transfer stage this is in hours using an average network speed of 10Mbps.
<sup>c</sup> Complexity for CPU hours is given in terms of classical computation time complexity; complexity for data transfer is shown in terms of how the size of the to be transferred data depends on the size of the input data.
<sup>\*</sup> This time is derived from the transfer of partial derivatives only, because for an association analysis with relatively few phenotypes it is not necessary to transfer encoded data.
$n_i$ - number of individuals in the study; $n_p$ - number of phenotypes of interest; $n_t$ - number of tests (genetic variants); $N_s$ - number of sites in the meta-analysis. In standard analysis $n_i \ll n_p$ and $n_i \ll n_t$

This drastic increase in performance is made possible through the shift of the computationally most expensive regression operation to the meta-analytical stage. For the meta-analytical stage, the HASE complexity is therefore slightly higher. However, it outperforms the classical meta-analysis using METAL (total computation time reduced 35 times), owing to the efficient implementation of our algorithm. Additionally, if HASE is only used for a high-dimensional association study of a single site, i.e. without subsequent meta-analysis with other sites, the computation time would be reduced 1400 times due to the removal of redundant calculations (for details see **Online Methods).**

### Application to real data

We used HASE to perform a high-dimensional association study in 4,034 individuals from the population-based Rotterdam Study. In this proof of principle study, we relate 8,723,231 million imputed genetic variants to 1,534,602 million brain magnetic resonance imaging (MRI) voxel densities (see **Online Methods).** The analysis was performed on a small cluster of 100 CPUs and took 17 hours to complete.

To demonstrate the potential of such high-dimensional analyses, we screened all genetic association results for both hippocampi (7,030 voxels) and identified the voxel with the lowest p-value. The most significant association (rs77956314; p = 3 x 10^−9^) corresponded to a locus on chromosome 12q24 **(Figure 1)**, which was recently discovered in a genome-wide association study of hippocampal volume encompassing 30,717 participants**^13^**.

**Figure 1.**
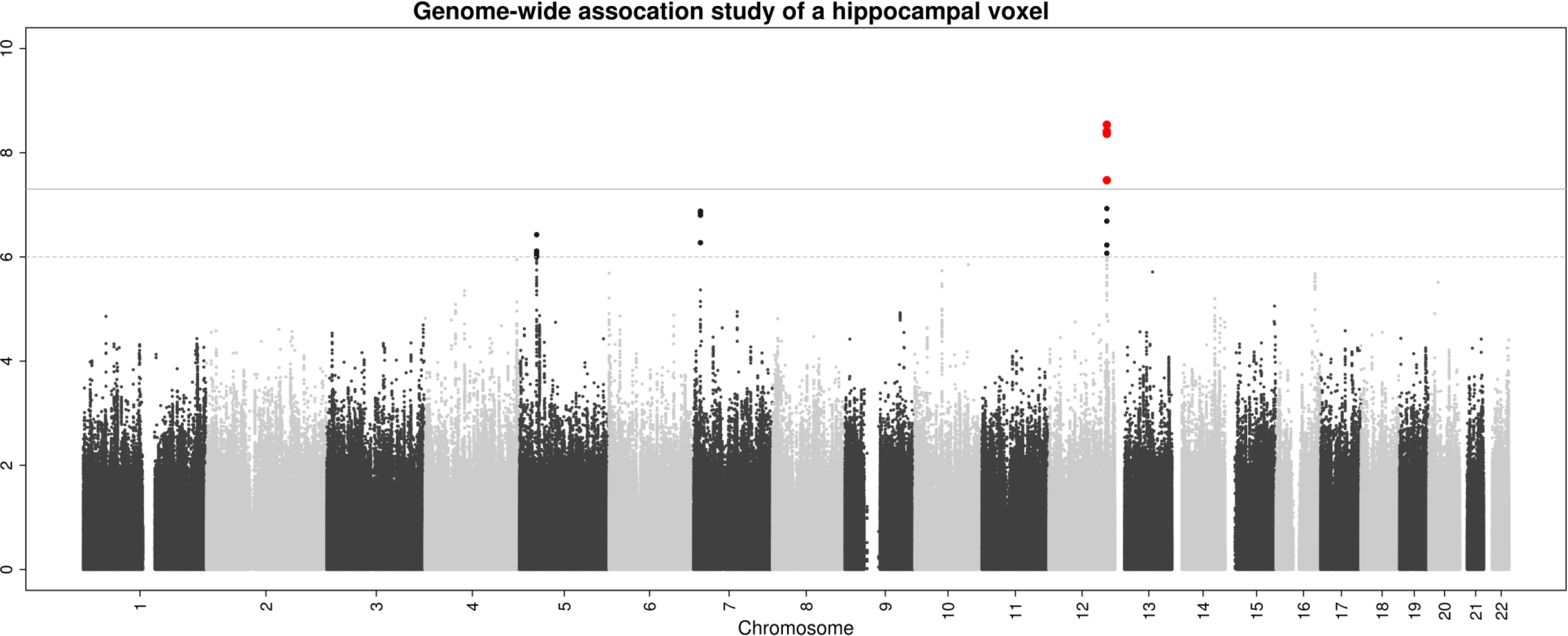
Manhattan plot of the hippocampus voxel with the most significant association after screening all 7030 hippocampal voxels. The most significant association (rs77956314; p = 3 ×10^−9^) corresponded to a previously identified locus on chromosome 12q24. Such voxel-wise hippocampus screening would take less than 8 hours on standard laptop.

Additionally, we performed the high-dimensional association studies separately in the three subcohorts of the Rotterdam Study and meta-analyzed the results using the HASE data reduction approach. It took on average 40 minutes for each subcohort to generate intermediate data for subsequent meta-analysis on a single CPU for all genetic variants and voxels. The meta-analysis was performed on the same cluster and took 17 hours to complete. Next, we compared the association results of the pooled analysis with the meta-analysis.**Figure 2** shows that the results are identical as it was predicted by theory (see **Online Methods).**

**Figure 2.**
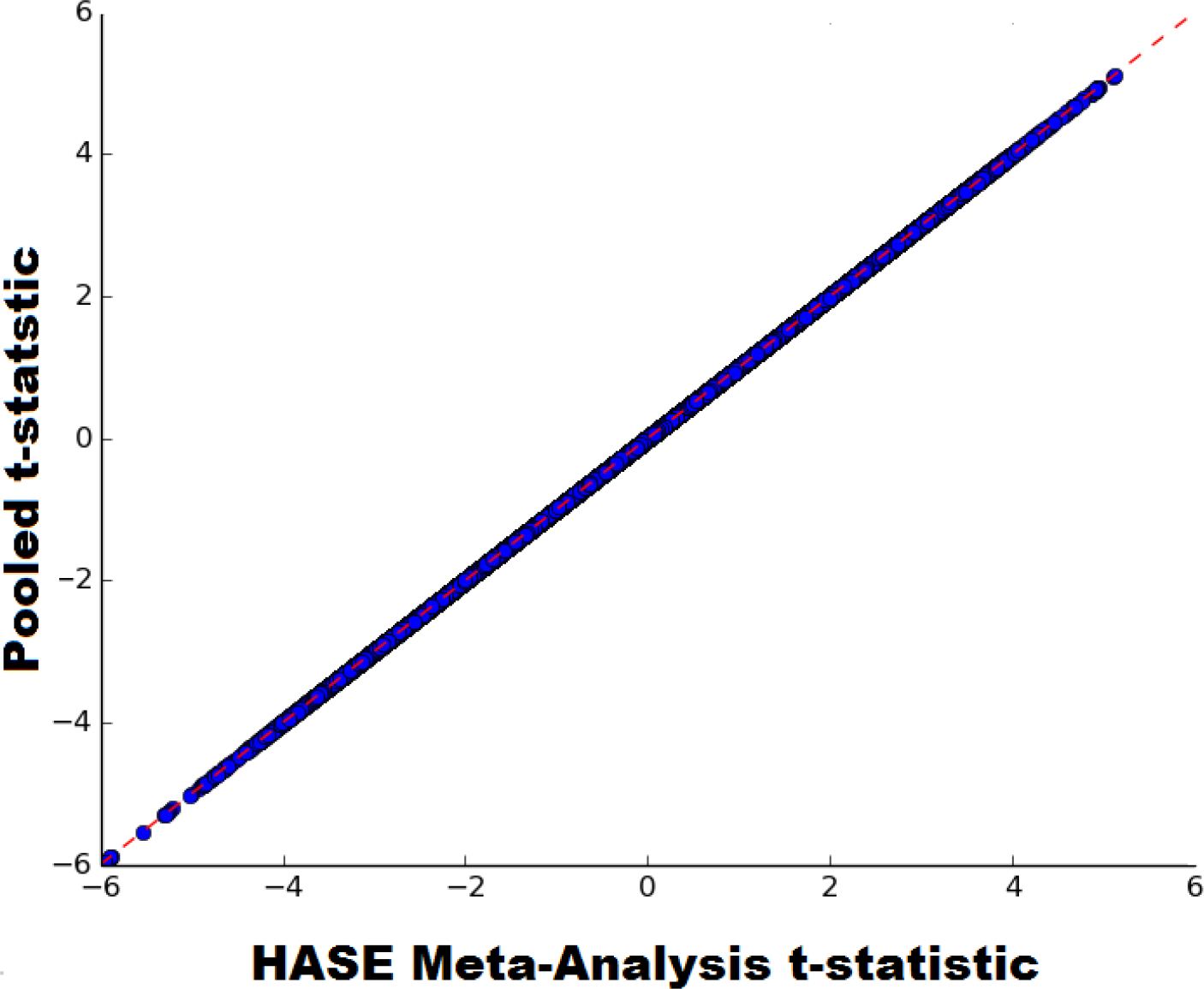
Correlation plot of voxel GWAS t-statistic estimated from pooled together data and voxel GWAS t-statistic estimated from meta-analysis of partial derivatives and encoded matrix. It took 40 min for single site to pre-compute data instead of 280 years to compute summary statistics; average data size to transfer was 17.5GBs instead of 300TBs.

## DISCUSSION

We describe a framework that allows for (i) computationally-efficient high-dimensional association studies within individual sites using standard computational infrastructure and (ii) facilitates the exchange of compact summary statistics for subsequent meta-analysis for association studies in a collaborative setting. Using HASE, we performed a genome-wide and brain-wide search for genetic influences on voxel densities, and illustrate both its feasibility and potential for driving scientific discoveries.

When using HASE, it is first necessary to convert the multi-dimensional data to a format that is optimized for fast reading and writing. This particular format, «hdf5», is not dependent on the architecture of the file system and can therefore be implemented on a wide range of hardware and software infrastructures. To facilitate this initial conversion step, we have built-in tools within the HASE framework for processing common file format of such big data. Furthermore, this is easily generalizable to other large data matrices in general and we foresee this initial conversion step not to form an obstacle for researchers to implement HASE.

In addition to the data format, a large improvement in efficiency comes from the reduced computational complexity. High-dimensional analyses contain many redundant calculations, which were removed in the HASE software. Also, we were able to further increase efficiency by simplifying the calculations to a set of matrix operations, which are computationally inexpensive, compared to conventional linear regression algorithms. Furthermore, the implementation of partial derivatives meta-analysis allowed us to greatly reduce the size of the summary statistics that need to be shared for performing a meta-analysis. Another advantage of this approach is that it only needs to calculate the partial derivatives for each site instead of the parameter estimates (i.e., beta coefficients and standard errors). This enabled us to develop within HASE a reduction approach that encodes data prior to exchange between sites, while yielding the exact same results after meta-analysis as if the original data were used. The encoding is performed such that tracing back to original data is impossible. This guarantees protection of participant privacy and circumvents restrictions on data sharing that are unfortunately common in many research institutions.

Alternative methods for solving the issues with high-dimensional data take one of two approaches. One approach is to reduce the dimensionality of the big datasets by summarizing the large amount of data into fewer variables^2^. Although this increases the speed, it comes at the price of losing valuable information, which these big data were primarily intended to capture. The second approach is to not perform a full analysis of all combinations of the big datasets, but instead make certain assumptions (e.g., a certain underlying pattern, or a lack of dependency on potential confounders) that allow for using statistical models that require less computing time. Again, this is a tradeoff between speed and accuracy, which is not necessary in the HASE framework, where computational efficiency is increased without introducing any approximations.

Unidimensional analyses of big data, such as genome-wide association studies, have already elucidated to some extent the genetic architecture of complex diseases and other traits of interest^1,14-16^, but much remains unknown. Cross-investigations between multiple big datasets potentially hold the key to fulfill the promise of big data in understanding of biology^7^. Using the HASE framework to perform high-dimensional association studies, this hypothesis is now testable.

**URLs.** Framework for efficient high-dimensional association analyses (HASE), https://github.com/roshchupkin/HASE/; description of the framework and protocol for metaanalysis, www.imagene.nl/HASE;

## Acknowledgments

The generation and management of GWAS genotype data for the Rotterdam Study are supported by the Netherlands Organization of Scientific Research NWO Investments (nr. 175.010.2005.011, 911-03-012). This study is funded by the Research Institute for Diseases in the Elderly (014-93-015; RIDE2), the Netherlands Genomics Initiative (NGI)/Netherlands Organization for Scientific Research (NWO) project nr. 050-060-810. The Rotterdam Study is funded by Erasmus Medical Center and Erasmus University, Rotterdam, Netherlands Organization for the Health Research and Development (ZonMw), the Research Institute for Diseases in the Elderly (RIDE), the Ministry of Education, Culture and Science, the Ministry for Health, Welfare and Sports, the European Commission (DG XII), and the Municipality of Rotterdam. MAI is supported by ZonMW grant number 916.13.054. HHHA is supported by the Van Leersum Grant of the Royal Netherlands Academy of Arts and Sciences. The Research is supported by the ImaGene programme of STW, The Society for Technical Scientific Research in The Netherlands.

## Author Contributions

GVR and HHHA jointly conceived of the study, participated in its design, performed the statistical analysis, interpreted the data, and drafted the manuscript. GVR additionally developed the software. MWV, AH, and CMvD acquired data and revised the manuscript critically for important intellectual content. MAI and WJN participated in its design, interpreted the data, and revised the manuscript critically for important intellectual content. All authors read and approved the final manuscript.

## Online Methods

### Hase

In high-dimensional associations analyses we test the following simple regression model:

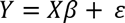

where **Y** is a **n_i_ × n_p_** matrix of phenotypes of interest, **n_i_** denotes the number of samples in the study, **n_p_** the number of phenotypes of interest, and ɛ denotes the residual effect. **X** is a three dimensional matrix **n_i_ × n_c_ × n_t_** of independent variables, with **n_c_** representing the number of covariates, such as the intercept, age, sex and, for example genotype as number of alleles, and **n_t_** the number of independent determinants.

In association analyses we are interested in estimating the p-value to test the null hypothesis that β=0. The p-values can be directly derived from the t-statistic of our test determinants. We will rewrite the classical equation for calculating t-statistics for our multi-dimensional matrices, which will lead to a simple matrix form solution for high-dimensional association analysis:

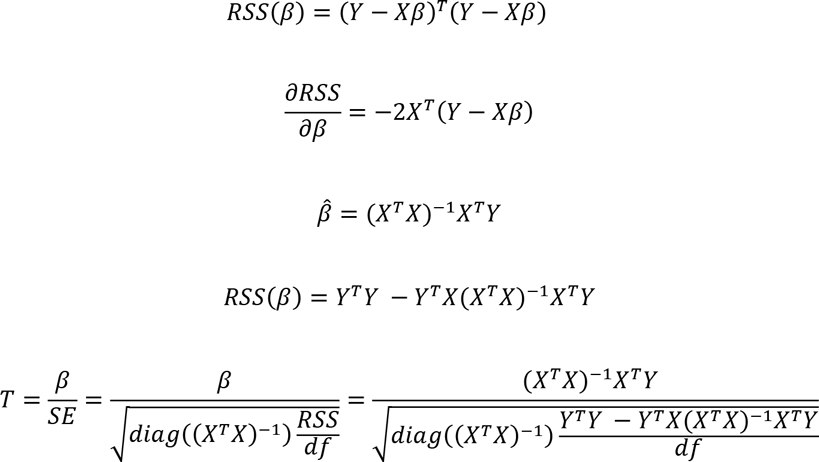

Where **T** is **n_p_ × n_c_ × n_t_** matrix of t-statistics and **df** is degree of freedom of our regression model. Let’s define A = X^T^X,B = X^T^Y and C = Y^T^Y, so that we can write our final equation for t-statistics:

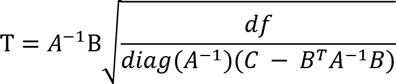

The result of this derivation is that, rather than computing all combinations of covariates and independent determinants, we only need to know three matrices: A, B and C, to calculate t-statistics and perform the full analysis. These results will be used in the section about meta-analysis.

The most computationally expensive operations here are the two multi-dimensional matrix multiplications (***A***^-1^**B**) and (***B^T^A^-^B***), where ***A*^-1^** is a three dimensional matrix **n_c_ × n_c_ × n_t_** and **B** is three dimensional matrix **n_c_ × n_p_ × n_t_**. Without knowledge of the data structure of these matrices, the simplest way to write the results of their multiplication would be to use Einstein’s notation for tensor multiplication:

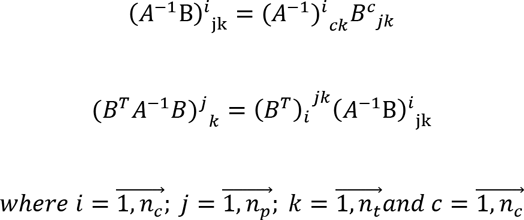

As you can see, the result is two matrices of **n_c_ × n_p_ × n_t_** and **n_p_ × n_t_** elements respectively. Despite the seemingly complex notation, the first matrix just represents the beta coefficients for all combinations of covariates (**n_c_** by **n_p_ × n_t_** combinations) and the second is fitting values of the dependent variable for every test (**n_p_ × n_t_** independent determinants).

However, insight into the data structure of **A** and **B** can dramatically reduce the computational burden and simplify operations. First of all, matrix **A** depends only on the covariates and number of determinants, making it unnecessary to compute it for every phenotype of interest, so we just need to calculate it once. Additionally, only the last covariate (i.e., the variable of interest) is different between tests, meaning that the (**n_p_ -1)×(n_p_ -1)xn_t_** part of matrix **A** remains constant during high-dimensional analyses. Matrix **B** consists of the dot product of every combination of the covariate and phenotype of interest. However, as we mentioned before, there are only (**n_t_ + n_c_ 1)** different covariates, and thus we can split matrix **B** in two low dimensional matrices: the first includes dot products of non-tested covariates - (**n_c_-1) × n_p_** elements; the second includes the dot products only of the tested covariates - **n_p_ × n_t_** elements. All this allows us to achieve large gain in computation efficiency and memory usage. In **Figure 1** we show a 2D schematic representation of these two matrices for standard genome association study with the covariates being an intercept, age, sex, and genotype. This example could be easily extrapolated to any linear regression model.

Applying the same splitting operation to ***B^T^*** it is possible to simplify tensor multiplication equation (**8**, **9**) to a low-dimensional matrix operation and rewrite the equation for t-statistics:

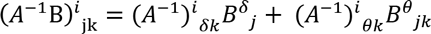

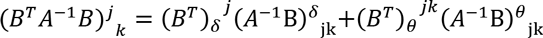

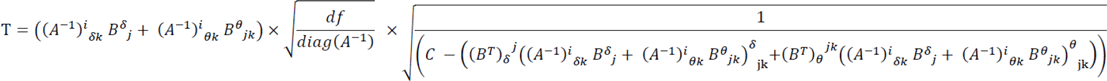

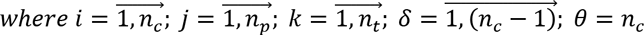

Then, to compute t-statistics for high-dimensional association analyses we just need to perform several matrix multiplications.

#### Meta-analysis

In classical meta-analysis, summary statistics such as beta coefficients and p-values are exchanged between sites. For 1.5 million phenotypes, this would yield around 400TB of data at each site, making data transfer to a centralized site impractical.

In the previous section we showed that, to compute all statistics for an association study, we just need to know the **A, B** and **C** matrices. As we demonstrated before^6^, by exchanging these matrices between sites, it is possible to gain the same statistical power as with a pooled analysis, without sharing individual participant data, because these matrices consist of aggregate data (**Figure 1).** However, in high-dimensional association analyses, matrix **B** grows very fast, particularly the part that depends on the number of determinants and phenotypes (**b_4_** in **Figure 1**).

If **Y** is a **n_i_ × n_p_** matrix of phenotypes of interest and **G** is a **n_i_ × n_t_** matrix of determinants which we want to test (e.g., a genotype matrix in GWAS), then **b_4_ = Y^T^** × **G.** These two matrices, **Y** and **G,** separately are not so large, but their product matrix has **n_p_ × n_t_** elements, which in a real application could be 10^6^ × 10^7^ =10^13^ elements and thus too large to share between sites. We propose to create a random **n_i_ × n_i_** nonsingular square matrix **F** and calculate its inverse matrix **F^-^**^1^. Then by definition **F × F^-1^=I,** where **I** is a **n_i_ × n_i_** elements identity matrix with ones on main diagonal and zeros elsewhere. Using this property, we can rewrite the equation for **b_4_**:

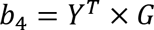

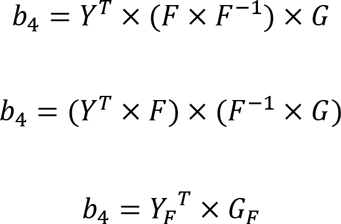

Therefore, instead of transferring TBs of intermediate statistics (**b_4_),** each side just needs to compute **A, C, Y_F_ and G_F_.**

Sharing just the encoded matrices does not provide information on individual participants and without knowing matrix **F** it is impossible to reconstruct the real data. However, it will be possible to calculate **b4,** perform a high-dimensional meta-analysis, and avoid problems with data transfer. Additionally, this method dramatically reduces computation time by shifting all complex computations to central site, where the HASE regression algorithm should be used to handle the association analysis in time efficient way.

**Figure 1.**
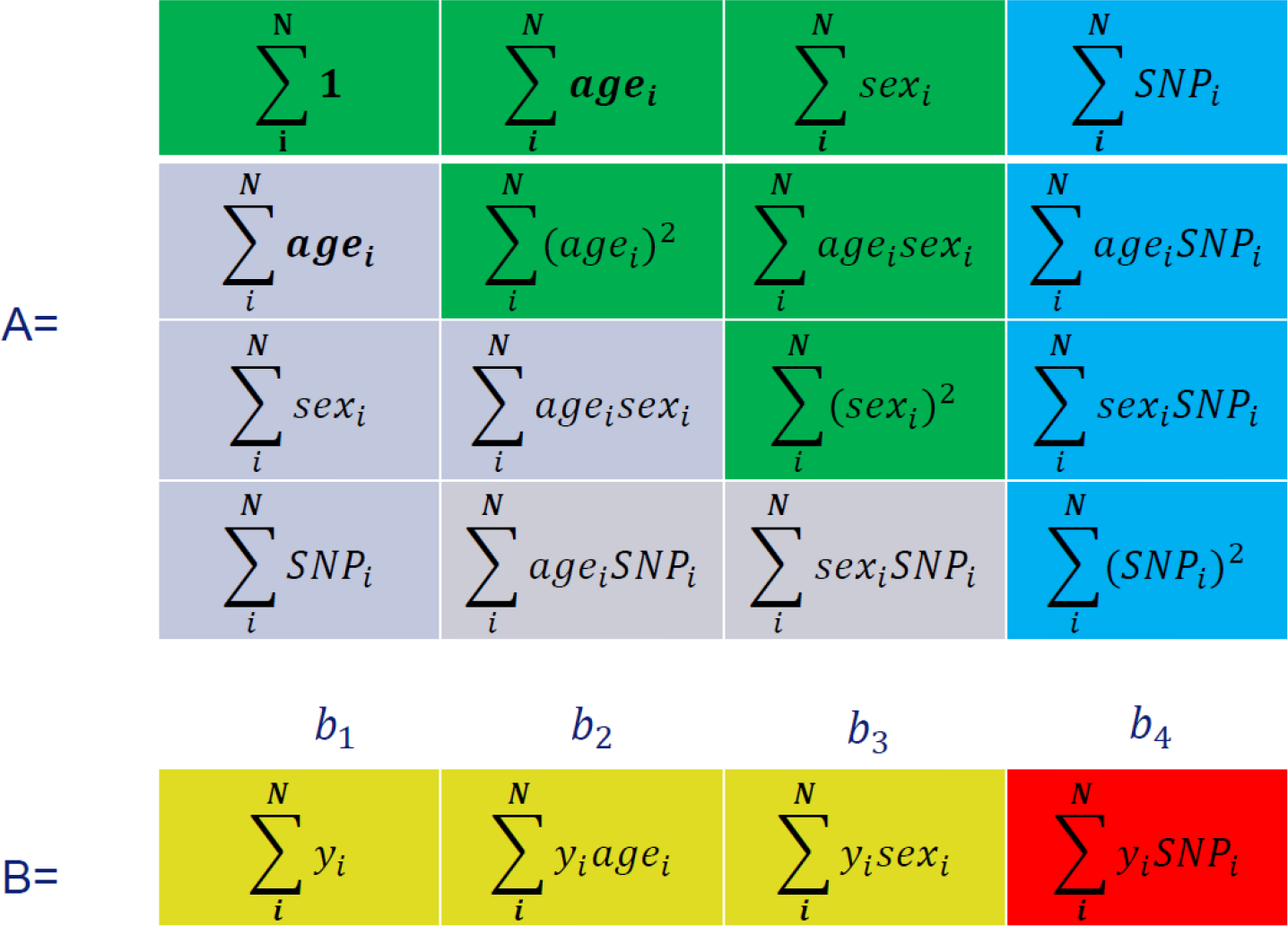
Explanation of the achieved speed reduction in HASE framework by removing redundant computations. In HASE multi-dimensional **A** and **B** matrices need to be calculated to perform GWAS studies. In the figure grey color means elements are parts of the matrix that are not necessary to calculate, as the **A** matrix is symmetric. The green color indicates elements that need to be calculated only once. Blue elements only have to be calculated for every SNP and yellow only for every phenotype. The red color indicates the most computationally expensive element, which needs to be calculated for every combination of phenotype and genotype. N denotes the number subjects in study.

### Supplementary Note

#### Study Population

The Rotterdam Study is an ongoing population-based cohort study in the Netherlands investigating diseases in the elderly and currently consists of 14,926 residents of Rotterdam who were aged 45 years or more at baseline [1,2]. The initial cohort was started in 1990 and expanded in 2000 and 2005. The whole population is subject to a set of multidisciplinary examinations every four years. MRI was implemented in 2005 and 5430 persons scanned until 2011 were eligible for this study. We excluded individuals with incomplete acquisitions, scans with artifacts hampering automated processing, participants with MRI-defined cortical infarcts, and subjects with dementia or stroke at the time of scanning. This resulted in a final study population of 4071 non-demented persons with information available on both genome-wide genotyping and MRI data. The Medical Ethics Committee of the Erasmus MC, University Medical Center Rotterdam and the review board of the Netherlands Ministry of Health, Welfare and Sports both approved the study. Informed consent was obtained from all subjects.

#### Imputation of genotypes

The Illumina 550K and 550K duo arrays were used for genotyping. Samples with low call rate (<97.5%), with excess autosomal heterozygosity (>0.336) or with sex-mismatch were excluded, as were outliers identified by the identity-by-state clustering analysis (outliers were defined as being >3 standard deviation (SD) from population mean or having identity-by-state probabilities >97%). A set of genotyped input SNPs with call rate >98%, MAF >0.001 and Hardy-Weinberg equilibrium (HWE) P-value > 10^−6^ was used for imputation. The Markov Chain Haplotyping (MACH) package version 1.0 software (Imputed to plus strand of NCBI build 37, 1000 Genomes phase I version 3) and minimac version 2012.8.6 were used for imputation.

#### MRI data

From August 2005 onwards, a dedicated 1.5 Tesla MRI scanner (GE Healthcare, Milwaukee, Wisconsin, USA) is operational in the Rotterdam Study research center in Ommoord. This scanner is operated by trained research technicians and all imaging data are collected according to standardized imaging protocols[2]. Brain MRI scans included a high-resolution 3D T1-weighted fast RF spoiled gradient recalled acquisition in steady state with an inversion recovery pre-pulse (FASTSPGR-IR) sequence with thin slices (voxel size<1mm^3^)[2]*Image processing*

Voxel based morphometry (VBM) was performed according to an optimized VBM protocol [3]. First, all T1-weighted images were segmented into supratentorial gray matter (GM), white matter (WM) and cerebrospinal fluid (CSF) using a previously described k-nearest neighbor (kNN) algorithm, which was trained on six manually labeled atlases [4]. FSL software [5] was used for VBM data processing. First, all GM density maps were non-linearly registered to the standard GM probability template. For this study we chose the ICBM MNI152 GM template (Montreal Neurological Institute) with a 1x1x1 mm^3^ voxel resolution. The MNI152 standard-space T1-weighted average structural template is derived from 152 structural images, which have been warped and averaged into the common MNI152 co-ordinate system after high-dimensional nonlinear registration.

A spatial modulation procedure was used to avoid differences in absolute GM volume due to the registration. This involved multiplying voxel density values by the Jacobian determinants estimated during spatial normalization. To gain more statistic power and decrease signal to noise ratio, all images were smoothed using a 3mm (FWHM 8mm) isotropic Gaussian kernel.

#### Statistical analysis

Linear regression models were fitted with voxel values of GM modulation density as the dependent variable and age, sex, and the number of minor alleles as independent variables. In total 1,534,602 voxels were processed.

